# Multi-omic and multi-view clustering algorithms: review and cancer benchmark

**DOI:** 10.1101/371120

**Authors:** Nimrod Rappoport, Ron Shamir

## Abstract

High throughput experimental methods developed in recent years have been used to collect large biomedical omics datasets. Clustering of such datasets has proven invaluable for biological and medical research, and helped reveal structure in data from several domains. Such analysis is often based on investigation of a single omic. The decreasing cost and development of additional high throughput methods now enable measurement of multi-omic data. Clustering multi-omic data has the potential to reveal further systems-level insights, but raises computational and biological challenges. Here we review algorithms for multi-omics clustering, and discuss key issues in applying these algorithms. Our review covers methods developed specifically for multi-omic data as well as generic multi-view methods developed in the machine learning community for joint clustering of multiple data types.

In addition, using cancer data from TCGA, we perform an extensive benchmark spanning ten different cancer types, providing the first systematic benchmark comparison of leading multi-omics and multiview clustering algorithms. The results highlight several key questions regarding the use of single-vs. multi-omics, the choice of clustering strategy, the power of generic multi-view methods and the use of approximated p-values for gauging solution quality. Due to the rapidly increasing use of multi-omics data, these issues may be important for future progress in the field.

## 1 Introduction

Deep sequencing and other high throughput methods measure a large number of molecular parameters in a single experiment. The measured parameters include DNA genome sequence [1], RNA expression [2, 3], DNA methylation [4] etc. Each such kind of data is termed “omic” (genomics, transcriptomics, methylomics, respectively). As costs decrease and technologies mature, larger and more diverse omic datasets are available. Computational methods are imperative for analyzing such data. One fundamental analysis is clustering - finding coherent groups of samples in the data, such that samples within a group are similar, and samples in different groups are dissimilar [5]. This analysis is often the first step done in data exploration. Clustering has many applications for biomedical research, such as discovering modules of co-regulated genes and finding subtypes of diseases in the context of precision medicine [6]. Clustering is a highly researched computational problem, investigated by multiple scientific communities, and a myriad algorithms exist for this task.

While clustering each omic separately reveals patterns in the data, integrative clustering using several omics for the same set of samples has the potential to expose more fine-tuned structures that are not revealed by examining only a single data type. For example, cancer subtypes can be defined based on both gene expression and DNA methylation together. Multi-omics clustering can also reduce the effect of experimental and biological noise in the data, and find structures that involve different cellular mechanisms.

A problem akin to multi-omics clustering was investigated independently by the machine learning community, and is termed “multi-view clustering” [7, 8]. Multi-view clustering algorithms can be used to perform clustering of multi-omic data. In the past, methods developed within the machine learning community have proven useful in the analysis of biomedical datasets. However, by and large, multi-view clustering have not penetrated bioinformatics yet.

In this paper, we review methods for multi-omics clustering, and benchmark them on real cancer data. The data source is TCGA (The Cancer Genome Atlas) [9] - a large multi-omic repository of data on thousands of cancer patients. We survey both multi-omics and multi-view methods, with the goal of exposing computational biologists to these algorithms. Throughout this review, we use the terms *view* and *multi-view* instead of omic and multi-omics in the context of Machine Learning algorithms.

Several recent reviews discussed multi-omics integration. [10], [11] and [12] review methods for multiomics integration, and [13] review multi-omics clustering for cancer application. These reviews do not include a benchmark, and do not focus on multi-view clustering. [14] reviews only dimension reduction multi-omics methods. To the best of our knowledge, [15] is the only benchmark performed for multi-omics clustering, but it does not include machine learning methods. Furthermore, we believe the methods tested in the benchmark do not represent the current state of the art for multi omics clustering. Finally, [7] is a thorough review of multi-view methods, directed to the Machine Learning community. It does not discuss algorithms developed by the bioinformatics community, and does not cover biological applications.

## 2 Review of multi-omics clustering methods

We divide the methods into several categories based on their algorithmic approach. *Early integration* is the most simple approach. It concatenates omic matrices to form a single matrix with features from multiple omics, and applies single-omic clustering algorithms on that matrix. In *late integration*, each omic is clustered separately and the clustering solutions are integrated to obtain a single clustering solution. Other approaches try to build a model that incorporates all omics, and are collectively termed *intermediate integration*. Those include: (1) methods that integrate sample similarities, (2) methods that use joint dimension reduction for the different omics datasets, and (3) methods that use statistical modeling of the data.

The categories we present here are not clear-cut, and some of the algorithms presented fit into more than one category. For example, iCluster [16] is an early integration approach that also uses probabilistic modeling to project the data to a lower dimension. The algorithms are described in the categories where we consider them to fit most.

Multi-omics clustering algorithms can also be distinguishable by the set of omics that they support. *General* algorithms support any kind of omics data, and are therefore easily extendible to novel future omics. *Omic specific* algorithms are tailored to a specific combination of data types, and can therefore utilize known biological relationships (e.g. the correlation between copy number and expression). A mixture of these two approaches is to perform feature learning in an omic specific way, but then cluster those features using general algorithms. For example, one can replace a gene expression omic with an omic that scores expression in cellular pathways, and thus take advantage of existing biological knowledge.

Throughout this review, we use the following notation: a multi-omic dataset contains *M* omics. *n* is the number of samples (or patients for medical datasets), *p_m_* is the number of features in the *m*’th omics, and *X^m^* is the *n* x *p_m_* matrix with measurements from the *m*’th omic. 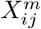 is therefore the value of the *j*’th feature for the *i*’th patient in the *m*’th omic. 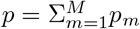 is the total number of features, and *X* is the *n* x *p* matrix formed by the concatenation of all *X^m^* matrices.

Figure 1 summarizes pictorially the different approaches to multi-omics clustering. A summary table of the methods reviewed here is given in Table 1.

**Figure 1:**
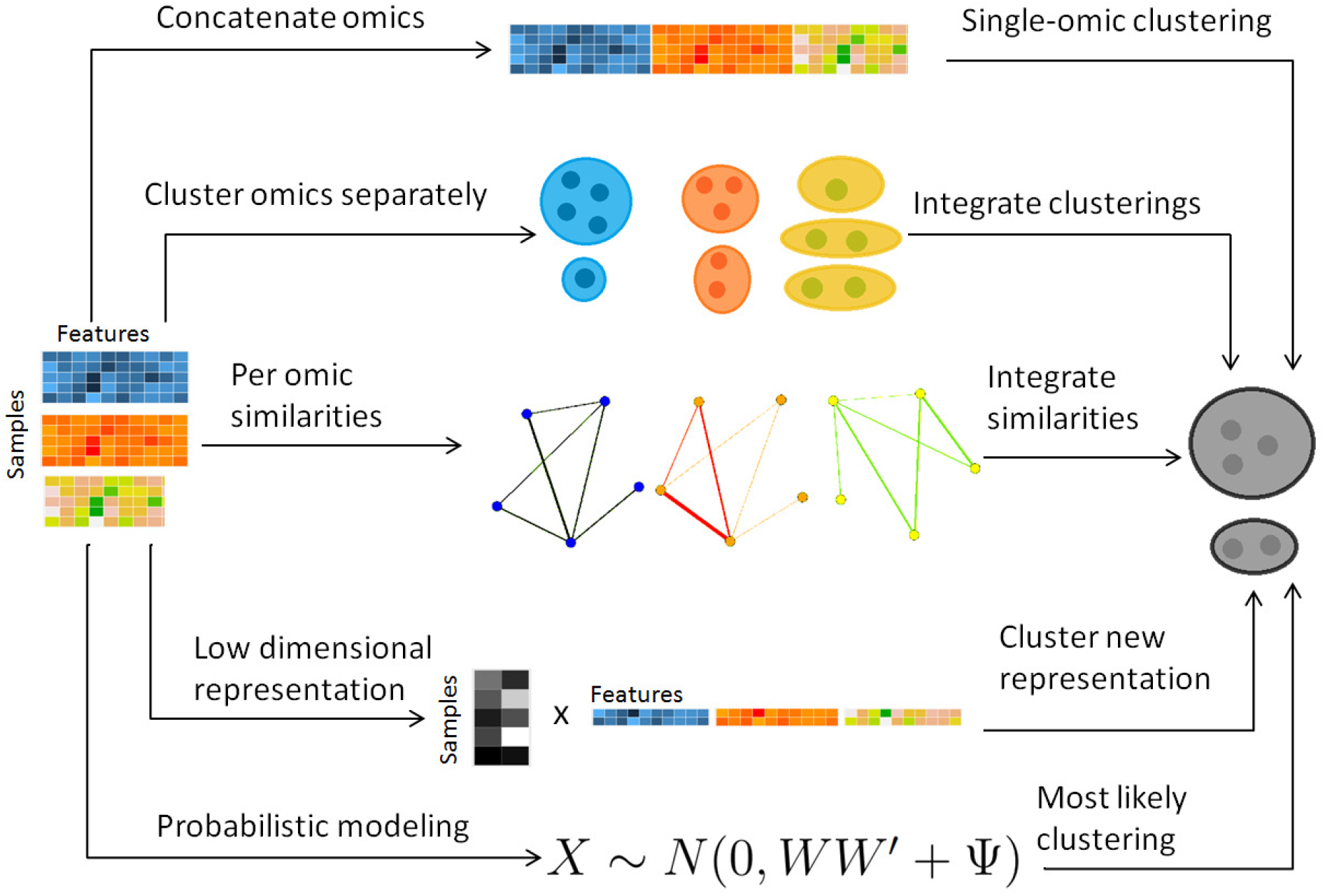
Overview of multi-omics clustering approaches.

**Table 1:** Multi-omic clustering methods. DR: dimension reduction; EM: Expectation maximization; MV: multi-view; NMF: Non-negative matrix factorization.^*•*^Methods included in the benchmark. Single-omic K-means and spectral clustering were also included in the benchmark.

| Method | Description | Refs. | Implementation |
| --- | --- | --- | --- |
| **Alternate optimization** |  |  |  |
| MV k-means, MV EM | Alternating k-means and EM. Each iteration is done w.r.t a different view | [17] | NA |
| **Early integration** |  |  |  |
| LRcluster<sup>•</sup> | Data originate from low rank matrix, omic data distributions modeled based on it | [18] | R |
| Structured sparsity | Linear transformation projects data into a cluster membership orthogonal matrix | [19] | Matlab |
| **Late integration** |  |  |  |
| COCA | Per omic clustering solutions integrated with hierarchical clustering | [20] | NA |
| Late fusion using latent models | Per omic clustering solutions integrated with PLSA | [21] | NA |
| PINS<sup>•</sup> | Integration of co-clustering patterns in different omics. The clusterings are based on perturbations to the data | [22] | R |
| **Similarity-based methods** |  |  |  |
| Spectral clustering generalizations | Generalizations of similarity based spectral clustering to multi-omics data | [23, 24, 25, 26] | Matlab |
| Spectral clustering with random walks | Generalizations of spectral clustering by random walks across similarity graphs | [? 27] | NA |
| SNF<sup>•</sup> | Integration of similarity networks by message passing | [28] | R, Matlab |
| rMKL-LPP<sup>•</sup> | DR using multiple kernel learning; similarities maintained in lower dimension | [29] | NA |
| **Dimension reduction** |  |  |  |
| General DR framework | General framework for integration with DR | [30] | NA |
| JIVE | Variation in data partitioned into joint and omic-specific | [31] | Matlab, R [32] |
| CCA<sup>•</sup> | DR to axes of max correlation between datasets. Generalizations: Bayesian, kernels, > 2 omics, sparse solutions, deep learning, count data | [33, 34, 35, 36, 37, 38, 39, 40, 41, 42, 43] | R, two omics [44], R, multiple omics |
| PLS | DR to axes of max covariance between datasets. Generalizations: kernels, > 2 omics, sparse solutions, partition into omic-specific and joint variation | [45, 46, 47, 48, 49, 50, 51, 52] | R, two omics, Matlab, multiple omics |
| MCIA | DR to axes of max covariance between multi-omic datasets | [53] | R |
| NMF generalizations<sup>•</sup> | DR using generalizations of NMF to multi-omic data | [54, 55, 56, 57, 58, 59, 60] | MultiNMF (Matlab) |
| Matrix tri-factorization | DR. Each omic describes the relationship between two entities | [61] | NA |
| Convex methods | DR with convex objective functions, allowing unique optimum and efficient computation | [62, 63, 18] | Matlab |
| Low-rank tensor MV clustering | Factorization based on low-rank tensors | [64] | Matlab |
| **Statistical methods** |  |  |  |
| iCluster/Plus/Bayes<sup>•</sup> | Data originate from low dimensional representation, which determines the distribution of the observed data | [16, 65, 66] | R |
| PARADIGM | Probabilistic model of cellular pathways using factor graphs | [67] | REST API |
| Disagreement between clusters | Methods based mainly on hierarchical Dirichlet processes; clustering in different omics need not agree | [68, 69, 70, 71, 72] | BCC (R) |
| Survival-based | Probabilistic model; patient survival data used in the clustering process | [73, 74] | SBC (R) |
| **Deep learning** |  |  |  |
| Deep learning methods | Neural networks used for integration. A variant of CCA, early integration and middle integration approaches | [36, 75, 76] | DeepCCA (Python) |

### 2.1 Alternate optimization

Early research for integration of two views was performed in [77]. This work improved classification accuracy for semi-supervised data with two views using an approach termed co-training, and inspired others to analyze multi-view data. One of the first attempts to perform multi-view clustering was [17]. In this work, EM and k-means, which are widely used single-omic clustering algorithms, were adapted for multi-view clustering. Both EM and k-means are iterative algorithms, where each iteration improves the objective function value. The suggested multi-view versions perform optimization in each iteration with respect to a different omic in an alternating manner. This approach loses theoretical guarantees for convergence, but was found to outperform algorithms that use each view separately, and also algorithms that cluster the concatenated matrix of the two views. Interestingly, [17] report improved results using the multi-view clustering algorithms on single-view datasets that were randomly split to simulate multi-view data. This was the first evidence for improved clustering using multiple views, and for the utility of a multi-view algorithm in clustering single-view data.

### 2.2 Early integration

Early integration is an approach that first concatenates all omic matrices, and then applies single-omic clustering algorithms on that concatenated matrix. It therefore enables the use of existing clustering algorithms. However, this approach has several drawbacks. First, without proper normalization, it may give more weight to omics with more features. Second, it does not consider the different distribution of data in the different omics. Finally, it increases the data dimension (the number of features), which is a challenge even in some single-omic datasets.

One way to handle the high dimension of the data is by using regularization, i.e., adding additional constraints to a problem to avoid overfitting [78]. Specifically, LASSO regularization creates models where the number of features with non-zero effect on the model is low [79], and regularization of the nuclear norm is often used to induce data sparsity. Indeed, LASSO regularization is used by iCluster [16] (reviewed in a later section), and LRACluster uses nuclear norm regularization (reviewed in this section). While any clustering algorithm can be applied using early integration, we highlight here algorithms that were specifically developed for this task.

LRACluster [18] uses a probabilistic model, where numeric, count and binary features have distributions determined by a latent representation of the samples Θ. For example, 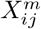 is distributed 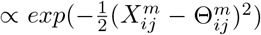, where Θ^*m*^ is of the same dimensions as *X^m^*. The latent representation matrix is encouraged to be of low rank, by adding a regularization on its nuclear norm. The objective function for the algorithm is −log(model’s likelihood) + *μ · |*Θ*|*_*_ where Θ is the concatenation of all Θ^*m*^ matrices, and *|*·*|*_*_ is the nuclear norm. This objective is convex and provides a global optimal solution, which is found using a fast gradient-ascent algorithm. Θ is subsequently clustered using k-means. This method was used to analyze pan-cancer TCGA data from eleven cancer types using four different omics, and to further find subtypes within these cancer types.

In [19], all omics are concatenated to a matrix *X* and the algorithm minimizes the following objective: 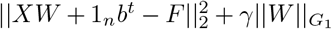. *W* is a *p* x *k* projection matrix, *F* is an *n* x *k* cluster indicator matrix such that *F^t^F* = *I_k_*, 1_*n*_ is a column vector of length *n* of 1’s, *b* is an intercept column vector of dimension *k* and *γ* is a scalar. The algorithm therefore seeks a linear transformation such the projected data are as close to a cluster indicator matrix as possible. That indicator matrix is subsequently used for clustering. The regularization term uses the *G*_1_ norm, which is the *l*_2_ norm for *W* entries associated with a specific cluster and view, summed over all views and clusters. Therefore, features that do not contribute to the structure of a cluster will be assigned with low coefficients in *W*.

### 2.3 Late integration

Late integration is another approach that allows to use existing single-omic clustering algorithms. First, each omic is clustered separately using a single-omic algorithm. Different algorithms can be used for each omic. Then, the different clusterings are integrated. The strength of late integration lies in that any clustering algorithm can be used for each omic. Algorithms that are known to work well on a particular omic can therefore be used, without having to create a model that unifies all of these algorithms. However, by utilizing only clustering solutions in the integration phase we can lose signals that are weak in each omic separately.

COCA [20] was applied to pan-cancer TCGA data, to investigate how tumors from different tissues cluster, and whether the obtained clusters match the tissue of origin. The algorithm first clusters each omic separately, such that the *m*’th omic has *c_m_* clusters. The clustering of sample *i* for omic *m* is encoded in a binary vector *υ_im_* of length *c_m_*, where *υ_im_*(*j*) = 1 if *i* belongs to cluster *j* and 0 otherwise. The concatenation of the *υ_im_* vectors across all omics results in a binary cluster indicator vector for sample *i*. The *n* x *c* binary matrix *B* of these indicator vectors, where 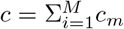, is used as input to consensus clustering [80] to obtain the final clustering of the samples. Alternatively, in [21] a model based on Probabilistic Latent Semantic Analysis [81] was proposed for clustering *B*.

PINS [22] integrates clusters by examining their connectivity matrices for the different omics. Each such matrix *S^m^* is a binary *n* x *n* matrix, where 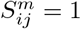 if patients *i* and *j* are clustered together in omic *m*, and 0 otherwise. These *S^m^* matrices are averaged to obtain a single connectivity matrix, which is then clustered using different methods based on whether the different *S^m^* matrices highly agree with each other or not. The obtained clusters are tested if they can be further split into smaller clusters. To determine the number of clusters for each omic and for the integrated clustering, perturbations are performed on the data by adding Gaussian noise to it, and the number of clusters is chosen such that the resulting clustering is robust to the perturbations.

Several methods for ensemble clustering were developed over the years, and are reviewed in [82]. While these were not originally developed for this purpose, they can be used for late multi-omics clustering as well.

### 2.4 Similarity-based methods

Similarity-based methods use similarities or distances between samples in order to cluster data. These methods compute the similarities between samples in each omic separately, and vary in the way these similarities are integrated. The integration step uses only similarity values. Since in current multi-omic datasets, the number of samples is much smaller than the number of features, these algorithms are usually faster than methods that consider all features while performing integration. However, in such methods it may be more difficult to interpret the output in terms of the original features. An additional advantage of similarity-based methods is that they can easily support diverse omic types, including categorical and ordinal data. Each omic only requires a definition of a similarity measure.

#### 2.4.1 Spectral clustering generalizations

Spectral clustering [83] is a widely used similarity-based method for clustering single-view data. The objective function for single-view spectral clustering is *max_U_ trace*(*U^t^LU*) s.t. *U^t^U* = *I*, where *L* is the Laplacian [84] of the similarity matrix of dimension *n* x *n*, and *U* is of dimension *n* x *k*, where *k* is the number of clusters in the data. Intuitively, it means that samples that are similar to one another have similar row vectors in *U*. This problem is solved by taking the *k* first eigenvectors of *L* (details vary between versions that use the normalized and the unnormalized graph Laplacian), and clustering them with a simple algorithm such as k-means. The spectral clustering objective was shown to be a relaxation of the discrete normalized cut in a graph, providing an intuitive explanation for the clustering. Several multi-view clustering algorithms are generalizations of spectral clustering.

An early extension to two views performs clustering by computing a new similarity matrix, using the two views’ similarities [23]. Denote by *W*_1_ and *W*_2_ the similarity matrices for the two views. Then the integrated similarity, *W*, is defined as *W*_1_*W*_2_. Spectral clustering is performed on the block matrix

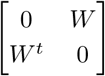

Note that each eigenvector for this matrix is of length 2*n*. Either half of the vector or an average of the two halves are used instead of the whole eigenvectors for clustering using k-means.

[24] generalizes spectral clustering for more than two views. Instead of finding a global *U* matrix, a matrix *U^m^* is defined for each omic. The optimization problem is:

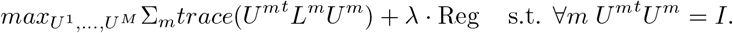

*L^m^* is the graph Laplacian for omic *m* and Reg is a regularization term equal to either 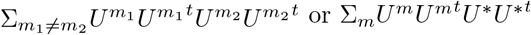 with the additional constraint that *U*^*^ is an *n* x *k* matrix such that *U ^*t^U ^*^* = *I*.

The first regularization allows each omic to have a different low rank *U^m^* representation, but requires that these representations are close to each other. The second regularization requires that the *U^m^* matrices are close to a consensus matrix *U^*^*. Each of the *U^m^* matrices, or *U^*^*, can then be used for clustering.

[25] uses a different formulation, which does not require a different *U^m^* for each omic, but instead uses the same *U* for all matrices. The following objective function is used:

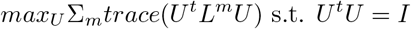

This is equivalent to performing spectral clustering on the Laplacian Σ*_m_L^m^*. The obtained clusters are then further improved in a greedy manner, by changing the assignment of samples to clusters, while looking directly at the discrete normalized cut objective, rather than the continuous spectral clustering objective.

[26] suggests a runtime improvement over [24]. Instead of looking at the similarity matrix for all the samples, a small set of “representative” vectors, termed salient points, are calculated by running k-means on the concatenation of all omics and selecting the cluster centers. A similarity matrix is then computed between these all samples in the data and their *s* nearest salient points. Denote this similarity matrix for the *m*’th omic by *W^m^*, and let *Z^m^* be its normalization such that rows sum to 1. These matrices are of dimension *n* x the number of salient points. Next, the matrices

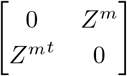

are given as input to an algorithm with the same objective as [25]. This way, similarities are not computed between all pairs of samples.

[?] views similarity matrices as networks, and examines random walks on these networks. Random walks define a stationary distribution on each network, which captures its similarity patterns [85]. Since that stationary distribution is less noisy than the original similarity measures, [?] uses them instead to integrate the networks. [27] also examines random walks on the networks, but argues that the stationary distribution in each network can still be noisy. Instead, the authors compute a consensus transition matrix, that has minimum total distance to the per-omic transition matrices and is of minimal rank.

#### 2.4.2 Similarity Network Fusion

SNF (Similarity Network Fusion) first constructs a similarity network for every omic separately [28]. In each such network, the nodes are samples, and the edge weights measure the sample similarity. The networks are then fused together using an iterative procedure based on message passing [86]. The similarity between samples is propagated between each node and its k nearest neighbors.

More formally, denote by *W*^(*m*)^ the similarity matrix for the *m*’th omic. Initially a transition probability matrix between all samples is defined by:

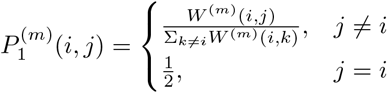

and a transition porbability matrix between nearest neighbors is defined by:

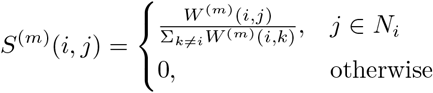

where *N_i_* are *i*’s k nearest neighbors in the input *X^m^* matrices. The *P* matrices are updated iteratively using message passing between the nearest neighbors: 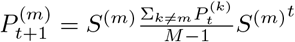 where 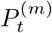 is the matrix for omic *m* at iteration *t*. This process converges to a single similarity network, summarizing the similarity between samples across all omics. This network is partitioned using spectral clustering.

In [28], SNF is used on gene expression, methylation and miRNA expression data for several cancer subtypes from TCGA. In addition to partitioning the graph to obtain cancer sutbypes, the authors show that the fused network can be used for other computational tasks. For example, they show how to fit Cox proportional hazards [87], a model that predicts prognosis of patients, with a constraint such that similar patients in the integrated network will have similar predicted prognosis.

#### 2.4.3 Multiple Kernel Learning

Kernel functions implicitly map samples to a high (possibly infinite) dimension, and can efficiently measure similarity between the samples in that dimension. Multiple kernel learning uses several kernels (similarity measures), and is often used in supervised analysis. [29] developed rMKL-LPP, which uses multiple kernel learning in unsupervised settings. The algorithm performs dimension reduction on the input omics such that similarities (defined using multiple kernels) between each sample and its nearest neighbors are maintained in low dimension. This representation is subsequently clustered with k-means. rMKL-LPP allows the use of diverse kernel functions, and even multiple kernels per omic. A regularization term is added to the optimization problem to avoid overfitting. The authors run the algorithm on five cancer types from TCGA, and show that using multiple kernels per omic improves the prognostic value of the clustering, and that regularization improves robustness.

### 2.5 Dimension reduction-based methods

Dimension reduction-based methods assume the data have an intrinsic low dimensional representation, with the dimension often corresponding to the number of clusters. The views that we observe are all transformations of that low dimensional data to a higher dimension, and the parameters for the transformation differ between views. This general formulation was proposed by [30], which suggest to minimize 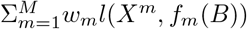, where *B* is a matrix of dimension *n* x *p*, *f_m_* are the parametrized transformations, and *w_m_* are weights for the different views, and *l* is a loss function. The work further provides an optimization algorithm when the *f_m_* transformations are given by matrix multiplication. That is, *f_m_*(*B*) = *BP^m^*, and *l* is the squared Frobenius norm applied to *X^m^ − BP^m^*. Once *B* is calculated, single-omic clustering algorithm can be applied to it. This general framework is widely used. Since the transformation is often assumed to be linear, many of the dimension reduction methods are based on matrix factorization. Dimension reduction methods work with real-valued data. Applying these methods to discrete binary or count data is technically possible but often inappropriate.

#### 2.5.1 JIVE

[31] assumes that the variation in each omic can be partitioned to a variation that is joint between all omics, and an omic-specific variation: *X^mt^* = *J^m^* + *A^m^* + *E^m^* where *E^m^* are error terms. Let *J* and *A* be the concatenated *J^m^* and *A^m^* matrices, respectively. The model assumes that *JA^t^* = 0, that is, the joint and omic specific variations are uncorrelated, and that *rank*(*J*) = *r* and *rank*(*A_i_*) = *r_i_* for each omic, so that the structure of each omic and the total joint variation are of low rank. In order for the weight of the different omics to be equal, the input omic matrices are normalized to have equal Frobenius norm. A penalty term is added to encourage variable sparsity. This method was applied to gene expression and miRNA data of Glioblastoma Multiforme brain tumors, and identified the joint variation between these omics.

#### 2.5.2 Correlation and covariance-based

Two of the most widely used dimension reduction methods are Canonical Correlation Analysis (CCA) [33] and Partial Least Squares (PLS) [45]. Given two omics *X*^1^ and *X*^2^, in CCA the goal is to find two projection vectors *u*^1^ and *u*^2^ of dimensions *p*_1_ and *p*_2_, such that the projected data has maximum correlation:

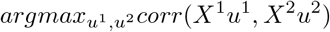

These projections are called the first canonical variates, and are the axis with maximal correlation between the omics. The k’th pair of canonical variates, 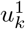 and 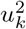 are found such that correlation between 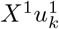 and 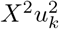 is maximal, given that the new pair is uncorrelated (that is, orthogonal) to the previous canonical variates. [88] proved and showed empirically that if the data originate from normal or log concave distributions, the canonical variates can be used to cluster the data. CCA was formulated in a probabilistic framework such that the optimization solutions are maximum likelihood estimates [89], and further extended to a Bayesian framework [34]. An additional expansion to perform CCA in high dimension is Kernel CCA [35]. A deeplearning based CCA method, DeepCCA, was recently developed [36]. Rather than maximize the correlation between linear projections of the data, the projections are taken to be functions of the data calculated using neural networks, and the optimization process optimizes the parameters for these networks.

Solving CCA requires inversion of the covariance matrix for the two omics. Omics data usually have a higher number of features than samples, and these matrices are therefore not invertible. To apply CCA to omics data, and to increase the interpretability of CCA’s results, sparsity regularization was added [37, 38].

CCA supports only two views. Several works extend it to more than two views, including MCCA [38] which maximizes the sum of pairwise correlations between projections and CCA-RLS [39]. [40] generalize CCA to tensors in order to support more than two views.

Another line of work on CCA, with high relevance for omics data, investigated relationships between the features while performing the dimension reduction. ssCCA (structure constrained sparse CCA) allows to incorporate into the model known relationships between features in one of the input omics, and force entries in the *u^i^* vector for that view to be close for similar features. This model has been developed by [41] and utilized microbiome’s phylogenies as the feature structure. Another model that considers relationship between features was developed in [42]. In this work, rather than defining similarities between features, they are partitioned into groups. Regularization is performed such that both irrelevant groups and irrelevant features within relevant groups are removed from the model. Finally, [43] extended CCA to support count data, which are common in biological datasets.

PLS also follows a linear dimension reduction model, but maximizes the covariance between the projections, rather than the correlation. More formally, given two omics *X*^1^ and *X*^2^, PLS computes a sequence of vectors 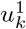 and 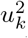 for *k* = 1, 2, *…* such that 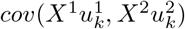 is maximal, given that 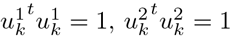, and 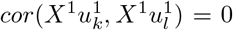 for *l* < *k*. That is, new projections are not correlated with previous ones. PLS can be applied to data with more features than samples even without sparsity constraints. A sparse solution is nonetheless desirable, and one was developed [46, 47]. O2-PLS increases the interpretability of PLS by partitioning the variation in the datasets into joint variation between them, and variations that are specific for each dataset and that are not correlated with one another [48]. While PLS and O2-PLS were originally developed for chemometrics, they were recently used for omics data as well [90, 91]. PLS was also extended to use the kernel framework [49], and a combined version of kernel PLS and O2 PLS was developed [50].

Like CCA, PLS was developed for two omics. MBPLS (Multi Block PLS) extends the model to more than two omics [92], and sMBPLS adds sparsity constraints. sMBPLS was developed specifically for omics data [51]. It looks for a linear combination of projections of non-gene-expression omics that has maximal correlation with a projection of gene expression omic. An extension of O2-PLS also exists for multi-view datasets [52].

An additional method that is based on maximizing covariance in low dimension is MCIA [53], an extension of co-inertia analysis to more than two omics [93]. It aims to find projections for all the omics such that the sum of squared covariances with a global variation axis is maximal: 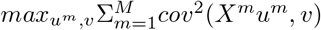. The projections of different omics can be used to evaluate the agreement between the different omics (the distance between projections reflects the level of disagreement between omics). Each of the projections can be used as a representation for clustering.

#### 2.5.3 Non-negative Matrix Factorization

Non-negative Matrix Factorization (NMF) assumes that the data have an intrinsic low dimensional non-negative representation, and that a nonnegative matrix projects it to the observed omic [94]. It is therefore only suitable for non-negative data. For a single omic, denote by *k* the low dimension. The formulation is *X ≈ WH*, where *X* is the *n* x *p* observed omic matrix, *W* is *n* x *k* and *H* is *k* x *p*. The objective function is 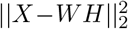, and it is minimized by updating *W* and *H* in an alternating manner, using multiplicative update rules, such that solutions remain non negative after each update [95]. The low dimension representation *W* can be clustered using a simple single-omic algorithm.

Several methods generalize this model to multi-omic data. MultiNMF [54] uses the following generalization: Each omic *X^m^* is factorized into *W^m^H^m^*. This model is equivalent to performing NMF on each omic separately. Integration between the omics is done by adding a constraint that the *W^m^* matrices are close to a “consensus” matrix *W*^*^. The objective function is therefore: 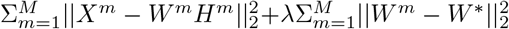. [55] generalizes this method to support weights for features’ and samples’ similarity. [56] extend MultiNMF by further requiring that the low dimensional representation *W*^*^ maintains similarities between samples (samples that are close in the original dimension must be close in *W*^*^). This approach combines factorization and similarity-based methods.

Joint NMF [57] uses a different formulation, where a sample has the same low dimensional representation for all omics: *X^m^ ≈ WH^m^*. Note that by writing *X* = *WH* where *X* and *H* are obtained by matrix concatenation, this model is equivalent to early integration. Joint NMF is not directly used for clustering. Rather, the data are reduced to a large dimension (*k* = 200) and high values in *W* and *H^m^* are used to associate samples and features with modules that are termed “md-modules”. The authors applied Joint NMF on miRNA, gene expression and methylation data from ovarian cancer patients, and showed that functional enrichment among features that are associated with md-modules that is more significant than the enrichment obtained in single-omic modules. In addition, patients in certain modules have significantly different prognosis compared to the rest of the patients. Much like [56] extends multiNMF, [58] extends Joint NMF such that similarities in the original omics are maintained in lower dimension. [59] extends NMF to the case where different views can contain different samples, but constrains certain samples from different views to belong to the same cluster based on prior knowledge. Finally, PVC [60] performs partial multi-view clustering. In this setting, not all samples necessarily have measurements for all views.

#### 2.5.4 Matrix tri-factorization

An alternative factorization approach presented in [61] is tri-matrix factorization. In this framework, each input omic is viewed as describing a relationship between two entities, which are its rows and columns. For example, in a dataset with two omics, gene expression and DNA methylation of patients, there are three entities which are the patients, the genes and the CpG loci. The gene expression matrix describes a relationship between patients and genes, while the methylation matrix describes a relationship between patients and CpG loci.

Each omic matrix *R_ij_* of dimension *n_i_* x *n_j_* that describes the relationship between entities *i* and *j* is factorized as 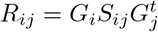, where *G_i_* and *G_j_* provide a low dimensional representation for entities *i* and *j* respectively and are of dimensions *n_i_* x *k_i_* and *n_j_* x *k_j_*, and *S_ij_* is an omic-specific matrix of dimension *k_i_* x *k_j_*. As in NMF, the *G_i_* matrices are non-negative. The same *G_i_* matrix is used in all omics with entity *i*, and in this way data integration is achieved. In the above example, both the gene expression and DNA methylation omics will use the same *G* matrix to represent patients, but different matrices to represent genes and CpG loci. In this model, an additional matrix describing the relationship between genes and CpGs could optionally be used. [61] adds constraints to the formulation that can encourage entities to have similar representations. This framework was applied to diverse problems in bioinformatics, including in supervised settings: It was used to perform gene function prediction [61], and for patient survival regression [96].

#### 2.5.5 Convex formulations

A drawback of most factorization-based methods is that their objective functions are not convex, and therefore optimization procedures do not necessarily reach a global optimum, and highly depend on initialization. One solution to this issue is by formulating dimension reduction as a convex problem. [62] relaxes CCA’s conditions and defines a convex variant of it. Performance was assessed on reducing noise in images, but the method can also be used for clustering. However, like CCA, the method only supports two views. [63] present a different convex formulation for dimension reduction, for the general factorization framework presented earlier, which minimizes 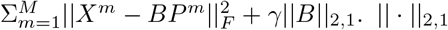 is the *l*_2,1_ norm, namely the sum of the Euclidean norms of the matrix rows. This relaxation therefore supports multiple views. LRAcluster [18] also uses matrix factorization and has a convex objective function.

#### 2.5.6 Tensor-based methods

A natural extension of factorization methods for multi-omic data is to use tensors, which are higher order matrices. One such method is developed in [64]. This method writes each omic matrix as *X^m^* = *Z^m^X^m^* + *E^m^*, *diag*(*Z^m^*) = 0, where *Z^m^* is an *n* x *n* matrix and *E^m^* are error matrices. The idea is that each sample in each omic can be represented as a linear combination of other samples (hence the *diag*(*Z^m^*) = 0 constraint), and that its representation in that base (*Z^m^*) can then be used for clustering. To integrate the different views, the different *Z^m^* matrices are merged to a 3^*rd*^-order tensor, *Z*. The objective function encourages *Z* to be sparse, and the *E^m^* error matrices to have a small norm.

### 2.6 Statistical methods

Statistical methods model the probabilistic distribution of the data. Some of these methods view samples as originating from different clusters, where each cluster defines a distribution for the data, while other methods do not explicitly use the cluster structure in the model. An advantage of the statistical approach is that it allows to include biological knowledge as part of the model when determining the distribution functions. This can be done either using Bayesian priors or by choosing probabilistic functions, e.g. using normal distribution for gene expression data. For most formulations, parameter estimation is computationally hard, and different heuristics are used. Several models under the Bayesian framework allow for samples to belong to different clusters in different omics.

#### 2.6.1 iCluster and iCluster+

iCluster [16] assumes that the data originate from a low dimension representation, which determines the cluster membership for each sample: *X^mt^* = *W^m^Z* + *ϵ^m^*, where *Z* is a *k* x *n* matrix, *W^m^* is an omic specific *p_m_* x *k* matrix, *k* is the number of clusters and *ϵ^m^* is a normally distributed noise matrix. This model resembles other dimension reduction models, but here the distribution of noise is made explicit. Under this model iCluster maximizes the likelihood of the observed data with an additional regularization for sparse *W^m^* matrices. Optimization is performed using an EM-like algorithm, and subsequently k-means is run on the lower dimension representation of the data *Z* to get the final clustering assignments. iCluster was applied to breast and lung cancer, using gene expression and copy number variations. iCluster was also recently used to cluster more than ten thousand tumors from 33 cancers in a pan-cancer analysis [97]. Note that by concatenating all *W^m^* matrices to a single *W* matrix, and rewriting the model as *X^t^* = *WZ* + *ϵ*, iCluster can be viewed as an early integration approach.

iCluster’s runtime grows fast with the number of features, and therefore feature selection is essential before using it [28]. [16] only use genes located on one or two chromosomes in their analysis.

Since iCluster’s model uses matrix multiplication, it requires real-values features. An extension called iCluster+ [65] includes different models for numeric, categorical and count data, but maintains the idea that data originate from a low dimension matrix *Z*. For categorical data, iCluster+ assumes the following model:

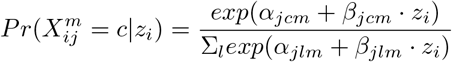

while for numeric data the model remains linear with normal error:

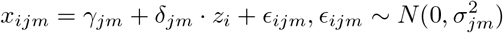

A regularization term encouraging sparse solution is added to the likelihood, and a Monte-Carlo Newton-Raphson algorithm is used to estimate parameters. The *Z* matrix is used as in iCluster for the clustering. The latest extension of iCluster, which builds on iCluster+, is iClusterBayes [66]. This method replaces the regularization in iCluster+ with full Bayesian regularization. This replacement results in faster execution, since the algorithm no longer needs to fine tune parameters for iCluster+’s regularization.

#### 2.6.2 PARADIGM

PARADIGM [67] is the most explicit approach to modeling cellular processes and the relations among different omics. For each sample and each cellular pathway, a factor graph that represents the state of different entities within that pathway is created. As a degenerate example, a pathway may include nodes representing the mRNA levels of each gene in that pathway, and nodes representing those genes’ copy number. Each node in the factor graph can be either activated, nominal or deactivated, and the factor graph structure defines a distribution over these activation levels. For example, if a gene has high copy number it is more likely that it will be highly expressed. However, if a repressor for that gene is highly expressed, that gene is more likely to be deactivated. PARADIGM infers the activity of non-measured cellular entities to maximize the likelihood of the factor graph, and outputs an activity score for each entity per patient. These scores are used to cluster cancer patients from several tissues.

PARADIGM’s model can be used for more than clustering. For example, PARADIGM-shift [98] predicts loss-of-function and gain-of-function mutations, by finding genes whose expression value as predicted based on upstream entities in the factor graph is different from their predicted expression value using downstream entities. PARADIGM relies heavily on known interactions, and requires specific modeling for each omic. It is also quite limited to the cellular level; For example, it is not clear how to incorporate into the model an omic describing the microbiome composition of each patient.

#### 2.6.3 Combining omic-specific and global clustering

All the methods discussed so far assume that there exists a consistent clustering structure across the different omics, and that analyzing the clusters in an integrative way will reveal this structure more accurately than analyzing each omic separately. However, this is not necessarily the case for biomedical datasets. For example, it is not clear that the methylation and expression profiles of cancer tumors really represent the same underlying cluster structure. Rather, it is possible that each omic represents a somewhat different cluster structure. Several methods take this view point using Bayesian statistics.

[68] defines a hierarchical Dirichlet process model, which supports clustering on two omics. Each sample can be either *fused* or *unfused*. Fused samples belong to the same cluster in both omics, while unfused samples can belong to different clusters in different omics. Patterns of fused and unfused samples reveal the concordance between the two datasets. This model is extended in PSDF [69] to include feature selection. [68] applies the model to cluster genes using gene expression and ChIP-chip data, while [69] clusters cancer patients using expression and copy number data.

In MDI [70] each sample can have different cluster assignments in different omics. However, a prior is given such that the stronger an association between two omics is, the more likely a sample will belong to the same cluster in these two omics. This association strength adjusts the prior clustering agreement between two omics. In addition to these priors, MDI’s model uses Dirichlet mixture model, and explicitly represents the distribution of the data within each cluster and omic. Since samples can belong to different clusters in different omics, no global clustering solution is returned by the algorithm. Instead, the algorithm outputs sets of samples that tend to belong to the same cluster.

A different Bayesian formulation is given by BCC [71]. Like MDI, BCC assumes a Dirichlet mixture model, where the data originate from a mixture of distributions. However, BCC does assume a global clustering solution, where each sample maps to a single cluster. Given that a sample belongs to a global cluster, its probability to belong to that cluster in each omic is high, but it can also belong to a different cluster in that omic. Parameters are estimated using Gibbs sampling [99]. BCC was used on gene expression, DNA methylation, miRNA expression and RPPA data for breast cancer from TCGA.

Like MDI and BCC, Clusternomics [72] uses a Dirichlet mixture model. Clusternomics suggests two different formulations. In the first, each omic has a different clustering solution, and the global clusters are represented as the Cartesian product of clusters from each omic. This approach does not perform integration of the multi-omic datasets. In the second formulation, global clusters are explicitly mapped to omic-specific clusters. That way, not all possible combinations of clusters from different omics are considered as global clusters.

#### 2.6.4 Survival-based clustering

One of the areas multi-omics clustering is widely used for is discovering disease subtypes. In this context, we may expect different disease subtypes to have a different prognosis, and this criterion is often used to assess clustering solutions. [73] develop a Bayesian model for multi-omics clustering that considers patient prognosis while clustering the data. Patients within a cluster have both similar feature distribution and similar prognosis. [74] also develop a probabilistic clustering method that considers survival, and that supports a large number of features compared to [73], which only uses a few dozen features. As the survival data are used as input to the model, it is not surprising that this approach gives clusters with more significantly different survival than other approaches. This was demonstrated on Glioblastoma Multiforme data by [73] and for data from several cancer types by [74], both from TCGA.

### 2.7 Deep multi-view methods

A recent development in machine learning is the advent of deep learning algorithms [100]. These algorithms use multi-layered neural networks to perform diverse computational tasks, and were found to improve performance in several fields such as image recognition [101] and text translation [102]. Neural networks and deep learning have also proven useful for multi-view applications [103], including unsupervised feature learning [36], [104]. Learned features can be used for clustering, as described earlier for DeepCCA. Deep learning is already used extensively for biomedical data analysis [105].

Recent deep learning uses for multi-omics data include [75] and [76]. [75] use an autoencoder, which is a deep learning method for dimension reduction. The authors ran it on RNA-seq, methylation and miRNA-seq data in order to cluster Hapatocellular Carcinoma patients. The architecture implements an early integration approach, concatenating the features from the different omics. The autoencoder outputs a representation for each patient. Features from this representation are tested for association with survival, and significantly associated features are used to cluster the patients. The clusters obtained have significantly different survival. This result is compared to a similar analysis using the original features, and features learned with PCA rather than autoencoders. However, the analysis in this work is not unsupervised, since the feature selection is based on patient survival.

[76] use a different approach. They analyze expression, methylation and miRNA ovarain cancer data using Deep Belief Networks [106] which explicitly consider the multi-omic strucutre of the data. The architecture contains separate hidden layers, each having inputs from one omic, followed by layers that receive input from all the single-omic hidden layers, thus integrating the different omics. A 3-dimensional representation over {0, 1} is learned for each patient, partitioning the patients into 2^3^ = 8 clusters. The clustering results are compared to k-means clustering on the concatenation of all omics, but not to other multi-omics clustering methods.

Deep learning algorithms usually require many samples and few features. They use a large number of parameters, which makes them prone to overfitting. Current multi-omic datasets have the opposite characteristics - they have many features and at least one order of magnitude less samples. The works presented here use only a few layers in their architectures to overcome this limitation, in comparison to the dozens of layers used by state-of-the-art architectures for imaging datasets. As the number of biomedical samples increases, deep multi-view learning algorithms might prove more beneficial for biomedical datasets.

## 3 Benchmark

In order to test the performance of multi-omics clustering methods, we compared nine algorithms on ten cancer types available from TCGA. We also compared the performance of the algorithms on each one of the single-omic datasets that make up the multi-omic datasets, for algorithms that are applicable to single-omic data. The nine algorithms were chosen to represent diverse approaches to multi-omics clustering. Three algorithms are early integration methods: LRAcluster, and k-means and spectral clustering on the omics concatenated into a single matrix. For similarity-based algorithms we used SNF and rMKL-LPP. For dimension reduction we used MCCA [38] and MultiNMF. We chose iClusterBayes as a statistical method, and PINS as a late integration approach.

The ten datasets contain cancer tumor multi-omics data, where each dataset is a different cancer type. All datasets contain three omics: gene expression, DNA methylation and miRNA expression. The number of patients range from 170 for AML to 621 for BIC. Full details on the datasets and cancer type acronyms appear in Supplementary File 2.

To assess the performance of a clustering solution, we used three metrics. First, we measured differential survival between the obtained clusters using the logrank test [107]. Using this test as a metric assumes that if clusters of patients have significantly different survival, they are different in a biologically meaningful way. Second, we tested for the enrichment of clinical labels in the clusters. We chose six clinical labels for which we tested enrichment: gender, age at diagnosis, pathologic T, pathologic M, pathologic N and pathologic stage. The four latter parameters are discrete pathological parameters, measuring the progression of the tumor (T), metastases (M) and cancer in lymph nodes (N), and the total progression (pathologic stage). Enrichment for discrete parameters was calculated using the *χ*^2^ test for independence, and for numeric parameters using Kruskal-Wallis test. Not all clinical parameters were available for all cancer types, so a total of 41 clinical parameters were available for testing. Finally, we recorded the runtime of each method. We did not consider in the assessment computational measures for clustering quality, such as heterogeneity, homogeneity or the silhouette score [108], since the different methods perform different normalization on the features (and some even perform feature selection). Full details about the survival and phenotype data appear in Supplementary File 2.

To derive a p-value for the logrank test, the *χ*^2^ test for independence, and the Kruskal-Wallis test, the statistic for these three tests is assumed to have *χ*^2^ distribution. However, for the logrank test and *χ*^2^ test this approximation is not accurate for small sample sizes and unbalanced cluster sizes, especially for large values of the test statistic (this was shown for example in [109] for the logrank test in the case of two clusters). Indeed, we encountered in our analysis cases where the approximation gave extreme p-values (< 10^−10^) for very small clusters (*n* = 3). Instead, we ran permutation tests for each clustering (where we permuted the cluster labels between samples) and used the test statistic to obtain an empirical p-value. The obtained empirical p-values were significantly different from the p-values returned by the *χ*^2^ approximation. In fact, for the logrank test on multi-omics data, the approximation-based p-values for 54 out of 89 clustering solutions (PINS crashed on BIC dataset, giving a total of 89 solutions) were not within their 95% confidence intervals constructed using the permutation test. This inaccuracy was exacerbated for small p-values - 32 out of the 35 significant (< 0.05) approximated p-values did not fall within their 95% confidence intervals. In all these cases, the p-value was higher (less significant) for the permutation-based computation. An extreme case is MCCA’s solution for KIRC dataset, where the p-value from the approximation was reported to equal 0, while the permutation test estimated it at 1.4e-4. In several cases, results that were significant according to the approximation were actually not significant according to the permutation tests. We observed a similar problem with the approximate p-values computed for the clinical parameters. The p-values we report here are therefore estimated using the permutation tests. More details on the permutation tests appear in Supplementary File 1. After permutation testing, the p-values for the clinical labels were corrected for multiple hypotheses (since several labels were tested) using Bonferroni correction for each cancer type and method at significance level 0.05. Results for the statistical analyses are in Supplementary File 3.

We applied all nine methods to the ten multi-omics datasets, and to the thirty single-omic matrices comprising them. The only exceptions were MCCA, which we could not apply to single-omic data, and PINS, which crashed consistently on all BIC datasets. All methods were run on a Windows machine, except for iCluster which was run on a Linux cluster utilizing up to 15 nodes in parallel. Details on hardware, data preprocessing and application of the methods appear in Supplementary File 1. Full clustering results appear in Supplementary File 4. All the processed raw data are available at http://acgt.cs.tau.ac.il/multi_omic_benchmark/download.html, and all software scripts used are available at https://github.com/Shamir-Lab/Multi-Omics-Cancer-Benchmark/.

Figure 2 depicts the performance of the benchmarked methods on the different cancer datasets, and Figures 3 and 4 summarize the performance for multi-omics data and for each single-omic separately across all cancer types. No algorithm consistently outperformed all others in either differential survival or enriched clinical parameters. With respect to survival, MCCA had the total best prognostic value (sum of −log10 p-values = 18.19), while MultiNMF was second (16.04) and rMKL-LPP third (14.18). The sum of p-values can be biased due to outliers, so we also counted the number of datasets for which a method’s solution obtains significantly different survival. These results are reported in Table 2. Here, with the exception of iClusterBayes, all methods that were developed for multi-omics or multi-view data had four cancer types with significantly different survival. These four cancer types are not identical for all the algorithms.

**Figure 2:**
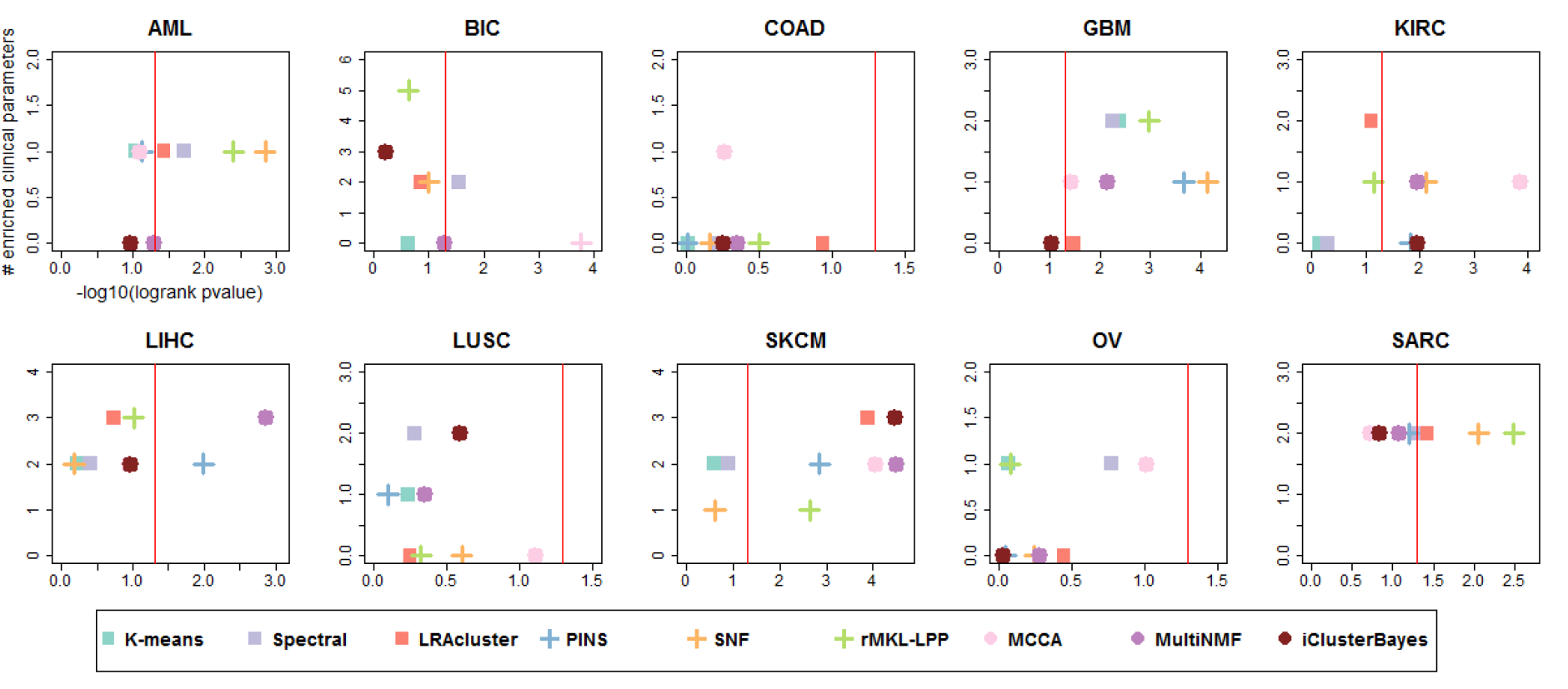
Performance of the algorithms on ten multi-omics cancer datasets. For each plot, the x-axis measures the differential survival between clusters (−log10 of logrank’s test p-value), and the y-axis is the number of clinical parameters enriched in the clusters. Red vertical lines indicate the threshold for significantly different survival (p-value ≤ 0.05)

**Figure 3:**
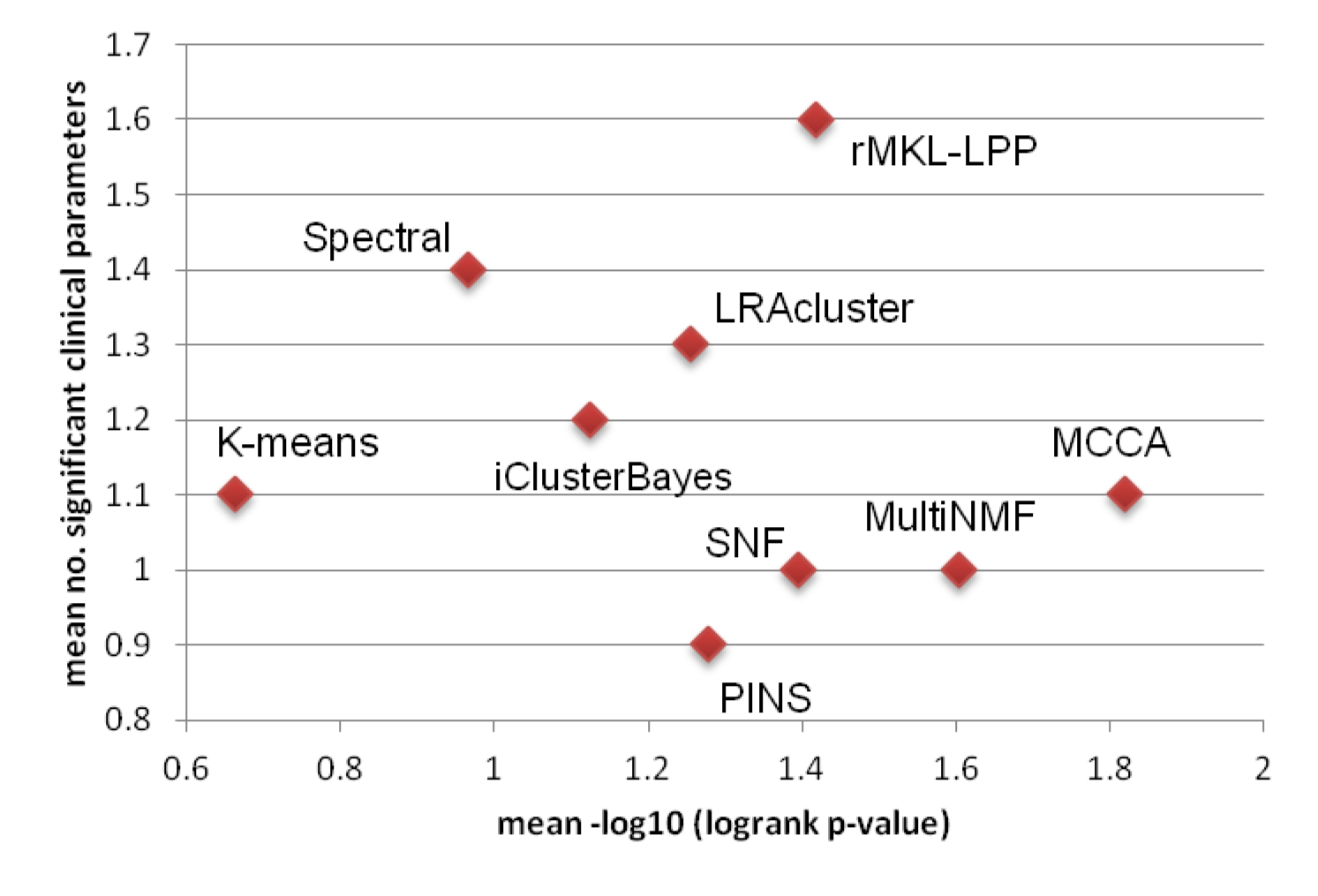
Mean performance of the algorithms on ten multi-omics cancer datasets. The x-axis measures the differential survival between clusters (mean −log10 of logrank’s test p-value), and the y-axis is the mean number of clinical parameters enriched in the clusters.

**Figure 4:**
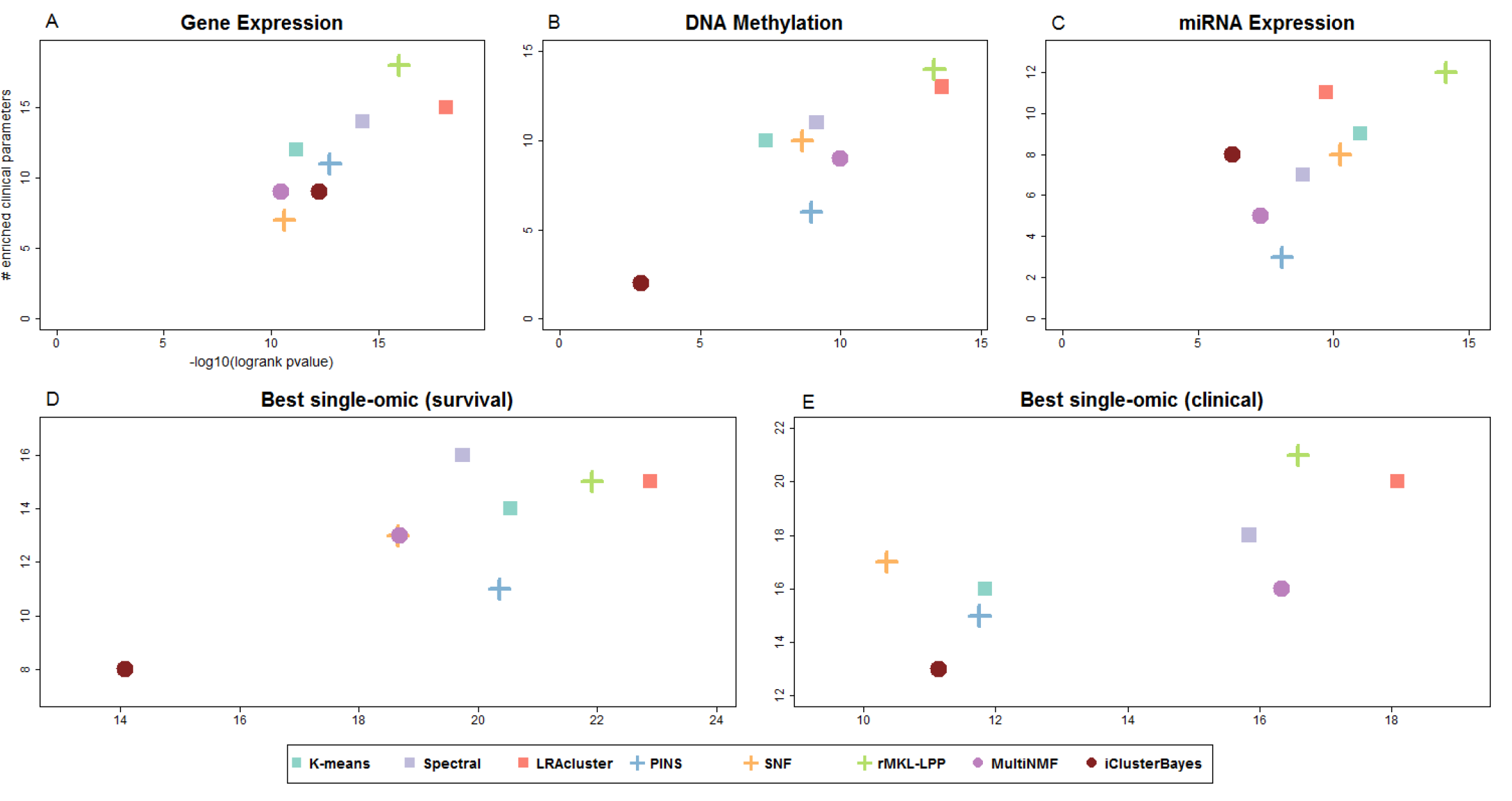
Summarized performance of the algorithms across ten cancer datasets. For each plot, the x-axis measures the total differential prognosis between clusters (sum across all datasets of −log10 of logrank’s test p-value), and the y-axis is the total number of clinical parameters enriched in the clusters across all cancer types. A-C: results for single-omic datasets. D: results when each method uses the single omic that achieves the highest significance in survival. E: same with respect to enrichment of clinical labels.

**Table 2:** Cancer types with significant results per algorithm. For each benchmarked algorithm, the number of cancer subtypes for which its clustering had significantly different prognosis (first row) and had at least one enriched clinical label (second row) are shown.

|  | K-means | Spectral | LRAcluster | PINS | SNF | rMKL-LPP | MCCA | MultiNMF | iClusterBayes |
| --- | --- | --- | --- | --- | --- | --- | --- | --- | --- |
| Significantly different survival | 1 | 3 | 4 | 4 | 4 | 4 | 4 | 4 | 2 |
| Significant clinical enrichment | 7 | 8 | 6 | 6 | 7 | 8 | 8 | 6 | 5 |

rMKL-LPP achieved the highest total number of significant clinical parameters, with 16 parameters. Spectral clustering came second with 14 and LRAcluster had 13. MCCA and MultiNMF, which had good results with respect to survival, had only 11 and 10 enriched parameters, respectively. rMKL-LPP did not outperform all other methods for all cancer types. For example, it had one enriched parameter for SKCM, while several other methods had two or three. We also considered the number of cancer types for which an algorithm had at least one enriched clinical label (Table 2). rMKL-LPP, spectral clustering and MCCA had enrichment in 8 cancer types, despite MCCA having a total of only 11 enriched parameters. Overall, rMKL-LPP outperformed all methods except MCCA and multiNMF with respect to both survival and clinical enrichment. MCCA and multiNMF had better prognostic value, but found less enriched clinical labels.

Each method determines the number of clusters for each dataset. These numbers are presented in Table 3. The numbers vary drastically among methods, from 2 or 3 (iCluster and MultiNMF) to more than 10 on average (MCCA). Both MCCA and rMKL-LPP partitioned the data into a relatively high number of clusters (average of 11.1 and 6.7 respectively), and both had good performance, which may indicate that clustering cancer patients into more clusters improves prognostic value and clinical significance. The higher number of clusters is controlled in the logrank and clinical enrichment tests by having more degrees of freedom for its *χ*^2^ statistic.

**Table 3:** Number of clusters chosen by the benchmarked algorithms on ten multi-omics cancer datasets. The right column is the average number of clusters across all cancer types.

|  | AML | BIC | COAD | GBM | KIRC | LIHC | LUSC | SKCM | OV | SARC | Means |
| --- | --- | --- | --- | --- | --- | --- | --- | --- | --- | --- | --- |
| K-means | 3 | 2 | 2 | 5 | 2 | 2 | 2 | 2 | 2 | 2 | 2.4 |
| Spectral | 9 | 3 | 2 | 5 | 2 | 2 | 2 | 2 | 4 | 2 | 3.3 |
| LRcluster | 6 | 10 | 6 | 13 | 7 | 15 | 14 | 6 | 9 | 10 | 9.6 |
| PINS | 4 | NA | 4 | 2 | 2 | 5 | 4 | 15 | 2 | 3 | 4.6 |
| SNF | 4 | 2 | 3 | 2 | 4 | 2 | 2 | 3 | 3 | 3 | 2.8 |
| rMKL-LPP | 6 | 7 | 6 | 6 | 11 | 6 | 6 | 7 | 6 | 6 | 6.7 |
| MCCA | 13 | 15 | 2 | 8 | 13 | 15 | 15 | 3 | 13 | 14 | 11.1 |
| MultiNMF | 2 | 2 | 2 | 3 | 2 | 3 | 2 | 2 | 2 | 2 | 2.2 |
| iClusterBayes | 2 | 3 | 2 | 2 | 2 | 3 | 2 | 2 | 2 | 2 | 2.2 |

The runtime of the different methods is reported in Table 4. Note that as mentioned earlier, iClusterBayes was run on a cluster, while the other methods were run on a desktop computer. All methods except for LRAcluster and iCluster took less than five minutes per dataset on average. LRAcluster and iClusterBayes took about 56 and 72 minutes per dataset, respectively.

**Table 4:** Runtime in seconds of the algorithms on ten multi-omics cancer datasets. The right column is the average runtime across all cancer types. *For iClusterBayes numbers are elapsed time on a multi-core platform.

|  | AML | BIC | COAD | GBM | KIRC | LIHC | LUSC | SKCM | OV | SARC | Means |
| --- | --- | --- | --- | --- | --- | --- | --- | --- | --- | --- | --- |
| K-means | 5 | 29 | 8 | 8 | 8 | 12 | 12 | 20 | 13 | 10 | 13 |
| Spectral | 3 | 8 | 3 | 3 | 3 | 5 | 5 | 6 | 4 | 4 | 4 |
| LRACluster | 935 | 11151 | 1442 | 1361 | 1007 | 4025 | 3459 | 6053 | 2361 | 2041 | 3384 |
| PINS | 41 | NA | 112 | 115 | 59 | 125 | 228 | 317 | 214 | 113 | 147 |
| SNF | 5 | 42 | 7 | 7 | 6 | 14 | 13 | 21 | 9 | 8 | 13 |
| rMKL-LPP | 222 | 192 | 205 | 221 | 191 | 255 | 213 | 333 | 263 | 238 | 233 |
| MCCA | 11 | 44 | 11 | 13 | 12 | 26 | 24 | 24 | 17 | 15 | 20 |
| MultiNMF | 19 | 51 | 25 | 19 | 17 | 35 | 27 | 45 | 21 | 23 | 28 |
| iClusterBayes* | 2628 | 7832 | 3213 | 2569 | 2756 | 5195 | 4682 | 6077 | 4057 | 3969 | 4298 |

Figure 4 also shows the performance of the benchmarked methods for single-omic data. While several methods had worse performance on single-omic datasets, some achieved better performance. For example, the highest number of enriched clinical parameters for both single and multi-omic datasets (18) was achieved by rMKL-LPP on gene expression. The gene expression solution also had better prognostic value than the multi-omic solution. LRAcluster on gene expression data had the most significant prognostic value across all single-omic and multi-omic experiments, except for MCCA on multi-omics data (sums of −log10 p-values are 18.15 and 18.19, respectively).

To further test how analysis of single-omic datasets compares to multi-omic datasets, we chose for each dataset and method the single omic that gave the best results for survival and clinical enrichment. In this analysis, rMKL-LPP had the highest total number of enriched clinical parameters (21), and the highest total survival significance was for LRAcluster (22.89). The runtime, number of clusters, and survival and clinical enrichment analysis for single-omic datasets appear in Supplementary Files 1 and 3. These results suggest that analysis of multi-omics data does not consistently provide better prognostic value and clinical significance compared to analysis of single-omic data alone, especially when different single-omics are used for each cancer types.

## 4 Discussion

We have reviewed methods for multi-omics and multi-view clustering. In our tests on ten cancer datasets, overall, rMKL-LPP performed best in terms of clinical enrichment, and outperformed all methods except MCCA and MultiNMF with respect to survival. The high performance of MCCA and MultiNMF is remarkable, as these are multi-view methods that were not specifically developed for omics data (though MCCA was applied to it).

Careful consideration should be given when applying multi-view clustering methods to multi-omic data, since these data have characteristics that multi-view methods do not necessarily consider. The most prominent of these characteristics is the large number of features relative to the number of samples. For example, CCA inverts the covariance matrix of each omic. This matrix is not invertible when there are more features than samples, and sparsity regularization is necessary. Another feature of multi-omic data is the dependencies between features in different omics, but several multi-view algorithms assume conditional independence of the omics given the clustering structure. This dependency is rarely considered, since it greatly increases the complexity of models. An additional characteristic of current omic data types is that due to cellular regulation, they have an intrinsic lower dimensional representation. The characteristic is utilized by many methods.

In our benchmark, single-omic data alone sometimes gave better results than multi-omics data. This was intensified when for each algorithm the “best” single-omic for each cancer type was chosen. These results question the current assumptions underlying multi-omics analysis in general and multi-omics clustering in particular.

Several approaches may lead to improved results for multi-omics analysis. First, methods that suggest different clusterings in different omics were developed and reviewed here, but were not included in the benchmark, since it is not clear how to compare algorithms that do not output a global clustering solution to those that do. These methods may be more sensitive to strong signals appearing in only some of the omics. Second, future algorithms can perform omic selection in the same manner that algorithms today perform feature selection. In the benchmark, we let each method choose a single-omic for each cancer type given the results of the analysis, which are usually not available for real data. Methods that filter omics with contradicting signals might obtain a clearer clustering. Finally, most methods for multi-omics clustering do not incorporate prior biological knowledge, and especially the relationship between omics. A notable exception is PARADIGM, which formulates both the relationships between different omics and between genes using known pathways. Other statistical methods also include some form of biological modeling by describing the distribution of the omics, and MDI tunes the similarity of clustering solutions in different omics based on the omics similarity. However, these methods do not model the biological relationship between omics. Methods that model such relations might benefit from additional biological knowledge, even without modeling whole pathways. For example, one can incorporate in a model the fact that promoter methylation is anti-correlated with gene expression. As far as we know, such methods were only developed for copy-number variation and gene expression data (e.g. [110]), and not in the context of clustering.

We detected large differences between the p-values derived from the *χ*^2^ approximation compared to the p-values derived from the permutation tests in the statistical tests we used. The differences were especially large due to the small sample size, small cluster sizes (in solutions with a high number of clusters) and due to a low number of events (high survival) for the logrank test. These p-values are used by single and multi-omic methods to assess their performance, and the logrank p-value is often the main argument for an algorithm’s merit. The large differences between the p-values question the validity of analyses that are based on the *χ*^2^ approximation, at least for TCGA data. Future work must use exact or permutation-based calculations of the p-value in datasets with similar characteristics to those used here for the benchmark.

The benchmark we performed is not without limitations. Gauging performance using patient survival is somewhat biased to known cancer subtypes, which may have been used in treatment decisions. Additionally, cancer subtypes that are biologically different may have similar survival. This is also true for enrichment of clinical parameters, although we attempted to choose parameters that would not lead to this bias. However, these measures are widely used for clustering assessment, including in the papers describing some of the benchmarked methods. Another limitation of the benchmark is that it only examines clustering, while some of the methods have additional goals and output. For example, in dimension reduction algorithms, the low dimensional data can be used to analyze features, and not only patients, e.g., by calculating axes of variation common to several omics. With respect to feature analysis, multi-omic algorithms can have an advantage over single-omic algorithms that we did not test. Finally, though we selected the parameters of each benchmarked method according to the guidelines given by the authors, judicious fine-tuning of the parameters may improve results.

## Acknowledgements

The results published here are based upon data generated by The Cancer Genome Atlas managed by the NCI and NHGRI. Information about TCGA can be found at http://cancergenome.nih.gov. We thank Nora K. Speicher for providing the rMKL-LPP tool and Ron Zeira for helpful comments.

## Funding

This research was supported in part by a grant from the United State - Israel Binational Science Foundation (BSF), Jerusalem, Israel and the United States National Science Foundation (NSF) and by the Bella Walter Memorial Fund of the Israel Cancer Association. N.R. was supported in part by a fellowship from the Edmond J. Safra Center for Bioinformatics at Tel-Aviv University.

